# Complete sequences of six Major Histocompatibility Complex haplotypes, including all the major *MHC class II* structures

**DOI:** 10.1101/2022.04.28.489875

**Authors:** Torsten Houwaart, Stephan Scholz, Nicholas R Pollock, William H. Palmer, Katherine M. Kichula, Daniel Strelow, Duyen B Le, Dana Belick, Tobias Lautwein, Thorsten Wachtmeister, Birgit Henrich, Karl Köhrer, Peter Parham, Lisbeth A Guethlein, Paul J Norman, Alexander T Dilthey

**Author notes:** These authors contributed equally (Torsten Houwaart, Stephan Scholz). These authors jointly supervised this work (Paul J Norman, Alexander T Dilthey).

## Abstract

Accurate and comprehensive immunogenetic reference panels are key to the successful implementation of population-scale immunogenomics. The 5Mbp Major Histocompatibility Complex (MHC) is the most polymorphic region of the human genome and associated with multiple immune-mediated diseases, transplant matching and therapy responses. Analysis of MHC genetic variation is severely complicated by complex patterns of sequence variation, linkage disequilibrium and a lack of fully resolved MHC reference haplotypes, increasing the risk of spurious findings on analyzing this medically important region. Integrating Illumina and ultra-long Nanopore sequencing as well as bespoke bioinformatics, we completed five of the alternative MHC reference haplotypes of the current (B38) build of the human reference genome and added one other. The six assembled MHC haplotypes encompass the DR1 and DR4 haplotype structures in addition to the previously completed DR2 and DR3, as well as six distinct classes of the structurally variable C4 region. Analysis of the assembled haplotypes showed that MHC class II sequence structures, including repeat element positions, are generally conserved within the DR haplotype supergroups, and that sequence diversity peaks in three regions around HLA-A, HLA-B+C, and the HLA class II genes. Demonstrating the potential for improved short-read analysis, the number of proper read pairs recruited to the MHC was found to be increased by 0.32% – 0.69% in a 1000 Genomes Project read re-mapping experiment with seven diverse samples. Furthermore, the assembled haplotypes can serve as references for the community and provide the basis of a structurally accurate genotyping graph of the complete MHC region.

## Introduction

The *Major Histocompatibility Complex* (*MHC*) is the most polymorphic region of the human genome, and is associated with more diseases, than any other region ^1-4^. The *MHC* spans 5Mbp of human chromosome 6 and encodes ∼165 proteins, as well as numerous *cis* and *trans*-acting factors ^5-8^. Over 40% of the encoded proteins are directly involved in immunity. Additional to HLA class I and II that control innate and adaptive immunity, are proteins that process the peptide substrates for presentation (e.g. TAP, tapasin, DM and DO), systemically acting complement components and cytokines (e.g. C4, TNFα), as well as transcription factors that promote or mediate immune responses (e.g. NFκB). Also encoded in the *MHC* are structural and developmental proteins (e.g. CDSN, NOTCH4), and other polymorphic molecules including MICA and HSP that act in stress-induced responses to infection. Most of the *MHC* region genes exhibit polymorphism and thus a strong potential to impact immune-mediated disease ^9^. Characterizing the *MHC* are intricate patterns of hyper-polymorphism, structural diversity and linkage disequilibrium (LD) ^10-12^. Such complexity arises through a dynamic evolutionary mechanism of natural selection and population demography ^13,14^. The dense sequence and structural diversity hinders attempts to comprehensively genotype *MHC* variation ^15^. The complex and insufficiently characterized patterns of LD, and to some extent the related functions of the encoded proteins, create additional challenges for fine mapping disease associations ^16,17^. For these reasons, it is of paramount importance to generate complete and accurate reference sequences that represent the extent of human genomic diversity in the *MHC* ^18,19^.

Within the *MHC*, most notable in their structural diversity and sequence divergence are the *MHC class II* and *C4* regions. The defining components of *MHC class II* include variable presence of the divergent *HLA-DRB3-5* genes and their LD with major *DRB1* variants ^20,21^, as well as clearly established hotspots of meiotic recombination ^22^. The defining characteristics of the *C4* region are gene duplication and resulting sequence homology, with up to four copies of the *C4* gene recorded per haplotype, as well as variable presence of a 19kb HERV insertion that affects expression of some alleles ^23^. Whereas imputation from dense SNP data has helped pinpoint some of the specific alleles associated with disease ^24^, low accuracy due to incompletely characterized sequence and LD patterns across populations, especially in the *C4* and *MHC class II* regions, can reduce the clinical utility of this approach ^25,26^. The most promising solutions for cataloguing *MHC* sequence complexity are graph-based approaches ^5^ that can represent all forms of genomic diversity without reference bias. The utility of these existing graph-based approaches beyond the classical *HLA* genes ^27-34^ remains limited because of a lack of fully resolved sequences that could be used to define a structurally accurate genotyping graph for *C4* and *MHC class II* and other genes in the *MHC* region.

Here we present and validate six fully resolved *MHC* sequences from homozygous cell lines as a reference for the community and as the basis for further methods development. We targeted five such cell lines (APD, DBB, MANN, QBL, SSTO) that were partially sequenced from BAC clones by the ‘*MHC Haplotype Project*’ ^35-38^ and further characterized from targeted short-read data ^9^. They form part of the current version of the human reference genome, GRCh38, being classified as alternative reference sequences (‘alt_ref’) for the *MHC* region ^39^. With our work, we increase the number of completely resolved *MHC* haplotypes of the cell lines present in GRCh38 from 2 to 7. In addition, we included one cell line, KAS116, that lacks *HLA-DRB3, -DRB4*, or *-DRB5* genes, thus representing one of two major *MHC class II* sequence structures currently not represented in GRCh38. Due to their importance for human health and difficulty of conventional genotyping, we focused our preliminary assessment of sequence features on the *MHC class II* and *C4* regions.

## Methods

### Culture of six *MHC*-homozygous cell lines

Six *MHC*-homozygous cell lines were cultured for fresh DNA extraction. We targeted five of the six cell lines that were partially sequenced as part of the ‘*MHC Haplotype Project*’ (the sixth, MCF, was unavailable at the time of study). In addition, we chose one cell line to represent the DR1 haplotype (Supplementary Table 1). All six cell lines were shown previously to be homozygous through the entire *MHC* region ^40,41^, and their *HLA class I* and *II* genotypes are shown in Supplementary Table 1. The cells investigated here are maintained by the European Collection of Cell Cultures (ECACC) and were purchased from Sigma-Aldrich, or obtained from the International Histocompatibility Working Group (IHWG) repository (http://www.ihwg.org/). Cells were cultured in RPMI 1640 (1X) media containing 100 mL FCS (20 %), 5 mL Penicillin/Streptomycin (1 %) and 5 mL Glutamine (2 mM). For initial seeding, cells were diluted to 3*10^5 cells/mL from cryopreserved stock. Cells were washed in PBS, pelleted beforehand by centrifugation (1200 rpm, 5 min.), and redistributed at 1*10^6 cells/mL. Cells were harvested at 1*10^7 cells/mL and pelleted.

### Single-molecule Nanopore sequencing of six *MHC*-homozygous cell lines

High molecular weight DNA was extracted following the protocol for ultra-long read nanopore sequencing ^42^. In summary, cells were lysed, digestion was performed using proteinase K, and DNA extraction using phenol/chloroform. Precipitated DNA was spooled onto a glass rod and washed in 70% Ethanol. Sequencing was carried out using the Oxford Nanopore sequencing platform, employing the MinION, GridION and PromethION devices following either the ultra-long protocol for library preparation ^42^ or the regular Oxford Nanopore Ligation Kits (SQK-LSK108/109). For ligation-based library preparation, ultra-high molecular weight DNA was sheared to a size of ∼75 kb using the Megaruptor 2 device and library preparation was performed following Oxford Nanopore’s protocol. Generated data for each cell line are summarized in Supplementary Table 2 and full details on the sequencing runs conducted are given in Supplementary Table 3.

### DNA sequencing (Illumina)

DNA was extracted for sequencing using the Qiagen Blood & Tissue Kit (Cat. No. 69506). Prior to library preparation 2500 ng of gDNA were sheared with Covaris ME220 (Covaris, Inc.) to a mean fragment size of 550bp. Library preparation was performed using the VAHTS Universal DNA Library Prep Kit for Illumina (Vazyme Biotech Co.; Ltd) according to the manufacturer’s protocol, without any amplification step, but with an additional size selection step after adapter ligation to remove smaller fragments. The library was quantified by qPCR by using KAPA library quantification kit (Roche Diagnostics Corporation) and a QuantStudio 3 (Thermo Fisher Scientific Inc.), and then sequenced using a HiSeq3000 system (Illumina Inc) with a read setup of 2×151 bp.

### Initial assembly of *MHC* sequences

Nanopore *MHC* reads were collected by alignment against previously assembled draft scaffolds of cell-line-specific contigs^9^, using minimap2 version 2.14-r892-dirty ^43^. Draft *MHC* assemblies were created from the Nanopore *MHC* reads using Canu ^44^ versions 1.7, 1.8, and 1.9, as well as Flye ^45^ version 2.6, empirically exploring the algorithms’ parameter spaces until a draft assembly containing a single *MHC* contig from a single algorithm had been obtained, which was then checked for structural consistency by long-read-to-assembly alignment and visual inspection. *MHC* draft assemblies were trimmed to “canonical” *MHC* coordinates by alignment against the GRCh38 *MHC* reference sequence (PGF) using nucmer ^46^ version 3.23, and polished in multiple iterations (see next section).

### Polishing *MHC* sequences

Polishing was carried out as an iterative process. Each iteration of polishing consisted of the following steps:

1. Nanopore polishing. Alignment of the Nanopore *MHC* reads against the input assembly sequence with minimap2^43^ and polishing using medaka 1.0.1[https://github.com/nanoporetech/medaka]; during the first round of polishing, medaka was used in ‘consensus’ mode; during the second and all subsequent rounds, medaka was used in ‘variant’ mode and the polishing was done by substituting homozygous variant calls into the assembly sequence according to the Medaka-generated VCF.
2. Illumina polishing. Whole-genome Illumina reads were aligned against a version of the primary (contains no ‘alt_refs’ or ‘decoys’) human reference genome GRCh38, in which the reference *MHC* sequence (PGF) was masked, and in which the output of step 1 was inserted as a separate contig, using BWA-MEM 0.7.15 ^47^. GATK ^48^ 4.1.4.1 variant calling was applied to the *MHC* contig, and a polished version of the *MHC* contig was produced by substituting homozygous variant calls into the assembly sequence according to the GATK-produced VCF.
3. Contig-based polishing. Cell-line-specific contigs ^9^ were aligned against the output of Step 2 and “high-quality” contig alignments were defined as alignments with “query cover” x “alignment identity” ≥ 0.99. The sequences of all high-quality contig alignments were substituted into the assembly sequence to obtain a polished assembly.

Polishing was carried out iteratively until manual inspection with IGV ^49^ and curation of the assemblies was indicative of high quality assembly. The number of polishing rounds per assembled *MHC* haplotype is listed in Supplementary Table 2.

After the last round of the iterative polishing process described above, a final round of a simple majority-based assembly improvement process was carried out: At each position in the assembled *MHC* haplotypes, the majority allele of the aligned *MHC* and Illumina *MHC* reads was determined using samtools 1.6 mpileup ^50^. If the Nanopore and Illumina majority alleles were in agreement and (i) accounted for more than 50% of aligned alleles at the considered position in their respective datasets; (ii) represented no INDEL; and (iii) disagreed with the allele carried by the assembly at the considered position, the assembly allele was replaced by the Nanopore+Illumina majority allele.

### Structural analysis

Multiple sequence alignments were computed and visualized using mauve ^51^.Pairwise sequence alignments were computed with nucmer (parameters --maxmatch --nosimplify --mincluster 300) version 3.1 ^46^ and visualized using mummerplot and gnuplot. For analysis of *MHC class II* sequences, analysis with mauve was found to be sensitive to selection of the ‘seed weight’ parameter. We empirically investigated multiple settings for ‘seed weight’ and settled on a value of 22 for visualization because it identified a reverse-complemented segment (see the Results section), the existence of which we verified with minimap 2. Note that, at ‘seed weight’ value 22, mauve generates non-syntenic alignments between different members of the *HLA-DRB* family for some haplotypes.

### Gene annotation

The generated *MHC* assemblies were annotated by comparison against the IMGT/HLA ^52^ database, comprising 14 genes and 25 pseudogenes, and the *MHC* components of the RefSeqGene (RSG) and RefSeq (RS) ^53^ databases, comprising 77 and 164 genes, respectively. For the genes and pseudogenes represented in IMGT/HLA, “genomic” sequences (representing full-length allelic variants of the included genes) were mapped ^43^ against the *MHC* assemblies. Alignments were filtered for complete alignments covering the query sequence in its entirety, and the highest-scoring alignment was selected to determine the start and end positions of the gene, for annotation purposes. If no complete alignment was found, no annotation was generated. For *MHC* genes not represented in IMGT, genomic reference sequences were extracted from RSG and, for genes not represented in RSG, from RS. Transcript structure was determined by projecting intron-exon boundaries from the query sequence of the selected highest scoring alignment onto the *MHC* assembly, and the deduced transcript was checked for translational consistency (presence of start and stop codons; absence of nonsense variants in coding regions). Except for transcripts previously shown ^9^ to encode incomplete gene products, no annotations were generated for which the implied transcript exhibited any translational inconsistencies. At the last step, annotation coordinates were converted from GFF into SQN format, using NCBI’s table2asn_GFF tool. Any transcripts failing the conversion process were removed.

### C4 genotyping

C4 polymorphism is characterized by three major factors of functional importance: gene copy number, antigenic determinants of the C4A and C4B isotypes, and presence (L) or absence (S) of a human endogenous retroviruses (HERV) insertion that reduces expression ^23,54^. C4 copy number and genotypes were determined by mapping the C4A sequence of GRCh38 ^39^, which contains the HERV element, to the assembled *MHC* sequences using minimap2 ^43^. The identified matches were classified with respect to HERV insertion and C4A/B status by (i) determining whether the corresponding alignment contained a deletion relative to the aligned C4A reference sequence between exons 9 and 10 (corresponding to HERV insertion status) and (ii) by determining whether the translated (i.e., amino acid) sequence of exon 26 contained the C4A/B-defining sequences ‘PCPVLD’ (C4A) or ‘LSPVIH’ (C4B).

### Repeat element annotation

Repeat elements were identified using RepeatMasker ^55^ version 4.1.2-p1. To obtain a common coordinate system for cross-haplotype display of repeat element positions, a multiple sequence alignment of the assembled *MHC* sequences was computed using mafft ^56^ version 7.490.

### Quantification of variants

In order to count variants of a given *MHC* haplotype sequence relative to the reference *MHC*, a pairwise sequence alignment was computed using minimap2 ^43^. The alignments corresponding to the mapped regions of the query sequence were projected onto the reference sequence, starting with the longest alignment. Each base of the query sequence was projected only once, even in the presence of overlapping supplementary alignments or secondary alignments. Small variants and INDELs were counted based on the projected pairwise alignments; structural variants (SVs) were defined as insertions or deletions of length ≥ 1000bp encoded by the CIGAR strings of the projected alignments, or as stretches along the query or reference sequences of more than 1000bp in length with no projected alignments. Of note, no separate analysis of inversions over and above the mauve-based analysis described above was performed during this step.

### Short-read mapping experiment

To investigate the effect on short-read mapping using the complete *MHC* assemblies presented here, (instead of the incomplete versions currently used in GRCh38), we performed a comparative mapping experiment. Seven samples were selected from the 1000 Genomes Project and their read data obtained from the resequencing effort by the New York Genome Center ^57^. Two genome references were created: (i) the 1000 Genomes Project ^58^ GRCh38-based reference genome with the HLA allele sequences removed and the ‘alt_ref’ *MHC* haplotypes retained; and (ii) a modified version in which the incomplete *MHC* haplotypes were substituted with the complete assemblies presented here, including KAS116 (which is not represented in GRCh38). In addition, a short *MHC* region ‘alt_ref’ contig (KI270758; 76,752bp) that we identified in GRCh38 and which maps to the KAS116 assembly was removed. Whole genome Illumina sequencing reads were aligned independently to each reference using BWA-MEM ^47^. Sample selection for the comparative mapping experiment was based on previously determined *HLA-DRB1* and *-DQB1* genotypes ^59,60^, targeting samples homozygous at these loci that matched the most (PGF) and least (APD) complete *MHC* references in GRCh38 (Supplementary Table 4)^35^. In addition, as we had sequenced KAS116 because it carries no *HLA-DRB3*/*4*/*5* genes, we selected *HLA-DRB1*08* and *HLA-DRB1*10* homozygous samples, which also carry no *HLA-DRB3*/*4*/*5* genes. For evaluating the comparative mapping experiment, we report (i) the number of reads aligning to any location in the utilized reference genome and (ii) the number of reads aligning to the included *MHC* sequences. For the MHC metric, only read alignments with the ‘proper pair’ flag set to 1 are considered, i.e. indicating the successful alignment of both reads of a read pair with correct read orientations and a plausible insert size.

## Results

### Assembly of six fully resolved *MHC* reference haplotypes

We integrated sequencing reads generated through ultra-long Nanopore, whole-genome Illumina and targeted Illumina ^9^ methods, together with tailored bioinformatics, to obtain high-quality, fully resolved assemblies of *MHC* haplotypes from five *MHC*-homozygous cell lines (Table 1a). These haplotypes (APD, DBB, MANN, QBL, SSTO) complete five of six unfinished versions present in the current build of the human reference genome (GRCh38) and complement the two completed haplotypes of PGF and COX already present ^35^. We also completed sequence and assembly of the *MHC* region from the KAS116 cell line using Nanopore and targeted Illumina sequence data only. The newly assembled haplotypes range in length from 4.90 to 5.05 Mbp (Figure 1). Compared with the GRCh38 versions of the same haplotypes, we resolved from 615 kbp (QBL) to 2.6 Mbp (APD) of additional DNA sequence. Compared to previously generated scaffolds ^9^, we resolved from 28kbp (APD) to 169 kbp (KAS116) of additional sequence (Supplementary Table 4). With the addition of KAS116, these four haplotypes represent the major *MHC class II* structural categories, DR1 (*DRB1* only; KAS116), DR2 (*DRB1*+*DRB5*; PGF), DR3 (*DRB1*+*DRB3*; APD, COX, QBL) and DR4 (*DRB1*+*DRB4*; DBB, MANN, SSTO).

**Table 1:** Sequenced cell lines and assembled haplotypes statistics. a) Summary of sequencing and assembly statistics. b) Summary of annotation and genomic variation. Structural variants are defined as insertions or deletions of more than 1000 bp in length. C4 genotypes are specified in a format including the C4 gene (“A” for C4A and “B” for C4B) and whether a HERV element insertion is present (“long”/L) or not (“short”/S).

| a. |  |  |  |  |  | b. |  |  |  |  |  |  |  |  |
| --- | --- | --- | --- | --- | --- | --- | --- | --- | --- | --- | --- | --- | --- | --- |
| Cell line | MHC class II type | MHC assembly (bp) | MHC class II (bp) | MHC read depth |  | GenBank ID | Cell line | Annotation |  |  | C4 genotype | Variants relative to PGF |  |  |
|  |  |  |  | Nanopore | Illumina (WG) |  |  | Gene | Pseudogenes | Transcripts |  | SNPs | INDELs | SVs |
| APD | DR3 | 4,929,194 | 149,490 | 31.65 | 16.26 | OK649231 | APD | 159 | 10 | 160 | AL,BL | 11,373 | 2,385 | 11 |
| DBB | DR4 | 5,048,539 | 260,755 | 20.03 | 12.91 | OK649232 | DBB | 155 | 10 | 154 | AL,BS | 13,310 | 2,595 | 19 |
| MANN | DR4 | 5,025,633 | 260,004 | 27.62 | 18.19 | OK649234 | MANN | 159 | 10 | 163 | BL | 12,777 | 2,280 | 17 |
| SSTO | DR4 | 5,046,870 | 258,832 | 22.43 | 15.04 | OK649236 | SSTO | 159 | 10 | 158 | AL,BL | 13,242 | 2,744 | 14 |
| KAS116 | DR1 | 4,911,495 | 156,116 | 85.87 | NA | OK649233 | KAS116 | 156 | 7 | 142 | AL,AL | 13,911 | 3,866 | 11 |
| QBL | DR3 | 4,904,958 | 149,415 | 17.81 | 12.86 | OK649235 | QBL | 158 | 10 | 160 | AL | 12,250 | 2,574 | 17 |
| COX | DR3 | 4,795,265 | 153,371 | NA | NA | GL000251.2 | COX | NA | NA | NA | BS | 12,278 | 2,021 | 16 |
| PGF | DR2 | 4,873,646 | 169,898 | NA | NA | chr6:28510120-33.480.575 (GRCh38) |  |  |  |  |  |  |  |  |

**Figure 1:**
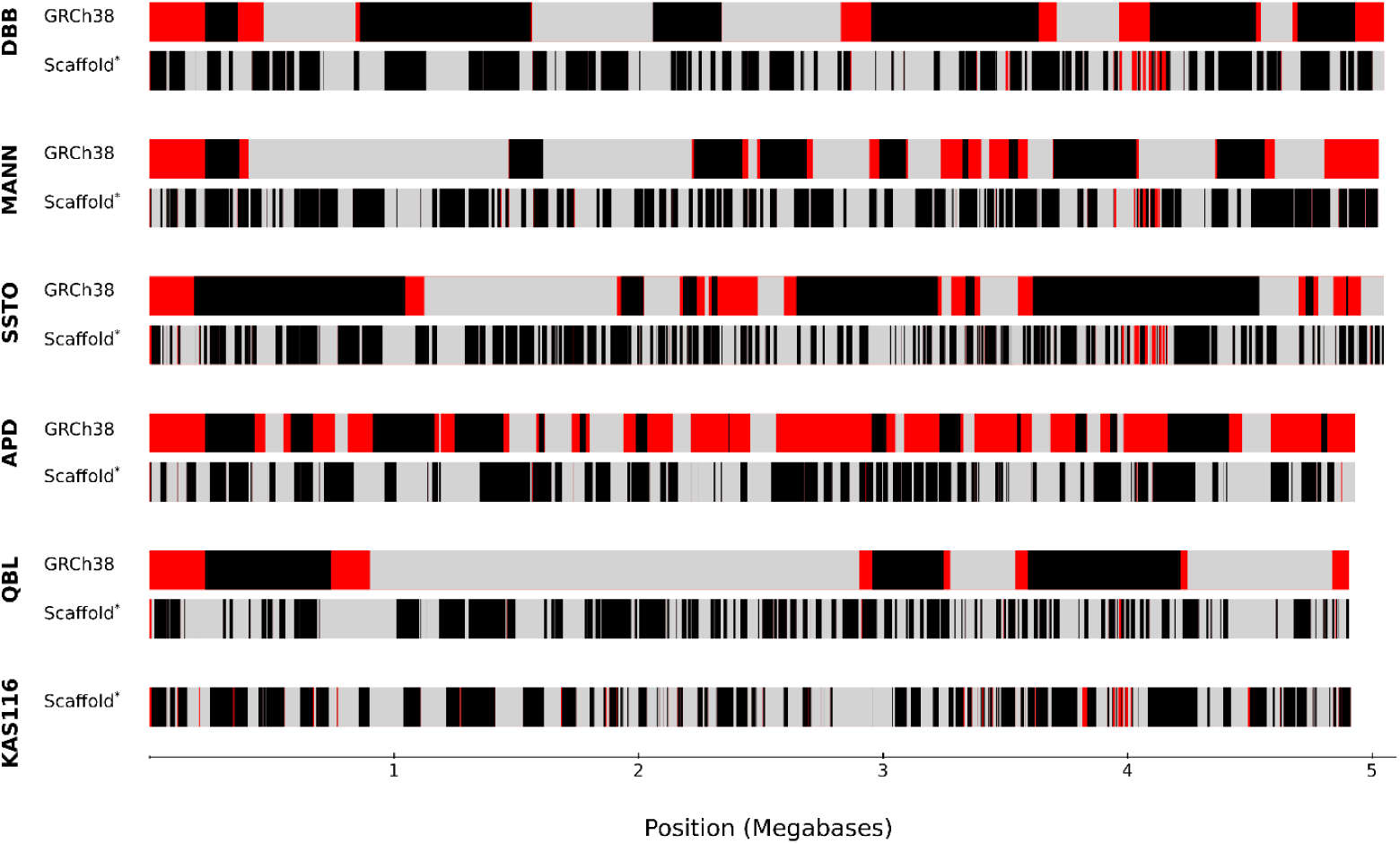
Contiguity and completeness increase over previous *MHC* assemblies. Shown for each of six haplotypes are comparisons to the assembly present in the current genome build (GRCh38: upper) and the short-read contigs scaffolds from ^9^ (SRC: lower). Regions absent from a previous assembly are coloured in red. Colour changes from black to grey or vice versa indicate distinct contigs that were not physically joined or separated by more than 5 undefined bases in the previous assemblies.

### Haplotype annotation and C4 genotypes

Using a semi-automated annotation approach (see Methods), we mapped the locations of 155 – 159 genes and their transcripts, and 7 – 10 pseudogenes onto the respective haplotypes (Table 1b). The annotations are included in the GenBank submissions of the sequences. Independently, we also determined the C4 genotypes (Table 2). The eight completed haplotypes represent six distinct classes of C4 status, C4-AL (QBL), AL,AL (KAS116), AL,BL (APD, PGF, SSTO) AL,BS (DBB), BL (MANN), BS (COX) (Supplementary Figure 1). Of note, we determined the C4 genotype of SSTO to be C4-AL,BL, in contrast to GRCh38, in which SSTO carries a C4-BS,BS structure.

**Table 2:** Comparative short-read mapping experiment. Comparison of whole-genome short-read mapping with and without completed MHC haplotypes.

| Sample ID | Sample *MHC class II* HLA types (homozygous) | Most similar to reference cell line | Mapping target | Reads aligned ('proper pairs') to the *MHC* | Total aligned reads (whole-genome) |
| --- | --- | --- | --- | --- | --- |
| NA20861 | DQB1*06:03 / DRB1*13:01 | APD (DQB1*06:03 / DRB1*13:01) | GRCh38 | 1,228,862 | 715,427,257 |
|  |  |  | GRCh38 + complete *MHC* | 1,235,188 | 715,427,774 |
|  |  |  | Difference | 0.51% | 0.00% |
| HG04206 | DQB1*06:03 /<br>DRB1*13:01 | APD<br>(DQB1*06:03 /<br>DRB1*13:01) | GRCh38 | 1,446,267 | 854,825,218 |
|  |  |  | GRCh38 +<br>complete<br>MHC | 1,452,415 | 854,825,865 |
|  |  |  | Difference | 0.43% | 0.00% |
| NA10847 | DQB1*05:01 /<br>DRB1*01:01 | KAS116<br>(DQB1*05:01 /<br>DRB1*01:01) | GRCh38 | 1,295,561 | 748,531,837 |
|  |  |  | GRCh38 +<br>complete<br>MHC | 1,304,461 | 748,532,066 |
|  |  |  | Difference | 0.69% | 0.00% |
| NA19755 | DQB1*04:01 /<br>DRB1*08:02 | KAS116 (no<br>DRB3/4/5) | GRCh38 | 1,192,504 | 702,458,873 |
|  |  |  | GRCh38 +<br>complete<br>MHC | 1,197,428 | 702,459,355 |
|  |  |  | Difference | 0.41% | 0.00% |
| HG02048 | DQB1*05:01 /<br>DRB1*10:01 | KAS116 (no<br>DRB3/4/5) | GRCh38 | 1,264,765 | 732,313,229 |
|  |  |  | GRCh38 +<br>complete<br>MHC | 1,270,691 | 732,313,730 |
|  |  |  | Difference | 0.47% | 0.00% |
| HG00135 | DQB1*06:02 /<br>DRB1*15:01 | PGF<br>(DQB1*06:02 /<br>DRB1*15:01) | GRCh38 | 314,505 | 212,840,164 |
|  |  |  | GRCh38 +<br>complete<br>MHC | 316,059 | 212,840,229 |
|  |  |  | Difference | 0.49% | 0.00% |
| NA11881 | DQB1*06:02 /<br>DRB1*15:01 | PGF<br>(DQB1*06:02 /<br>DRB1*15:01) | GRCh38 | 1,315,963 | 753,208,749 |
|  |  |  | GRCh38 +<br>complete<br>MHC | 1,320,137 | 753,209,300 |
|  |  |  | Difference | 0.32% | 0.00% |

### Large-scale *MHC* sequence structures and density of genetic variation

With exception of the *MHC class II* region (described below), no large-scale structural differences across the assembled haplotypes were identified, using either multiple sequence alignment (Supplementary Figure 2) or pairwise sequence alignment (Supplementary Figure 3) approaches. To assess the density of genetic variation across the *MHC*, we mapped the assemblies against the canonical PGF reference haplotype and quantified them in sliding windows. With the exception of the MHC *class II* region, the positioning (data not shown) and density (Supplementary Table 5) of repeat elements were generally conserved between the assembled haplotypes, with total interspersed repeats accounting for 51 – 52% of sequence content. Consistent with previous analyses ^9,61^, we observed three peaks having up to 50 SNPs / kbp, centred around the *HLA-A, HLA-B+C* and *HLA class II* genes respectively, and that structural diversity peaks in the *MHC class II* region (Figure 2). Overall numbers of variants are similar across the assemblies, with the exception of an increased INDEL count observed for KAS116 (3,866 compared to 2280 – 2744 for the other assemblies; Table 1b), possibly reflecting an increased INDEL error rate due to the fact that KAS116 was not polished with whole-genome Illumina sequencing data. Consistent with this possibility, the number of annotated complete transcripts was slightly reduced for KAS116 (142 compared to e.g. 154 for DBB), possibly reflecting an increased rate of apparent ‘missense’ mutations in transcripts induced by INDEL assembly errors. When treating COX as an assembly gold standard for the comparison with PGF, the observed increased count of INDELs for KAS116 would translate into one additional INDEL error approximately every 2,600 bp.

**Figure 2:**
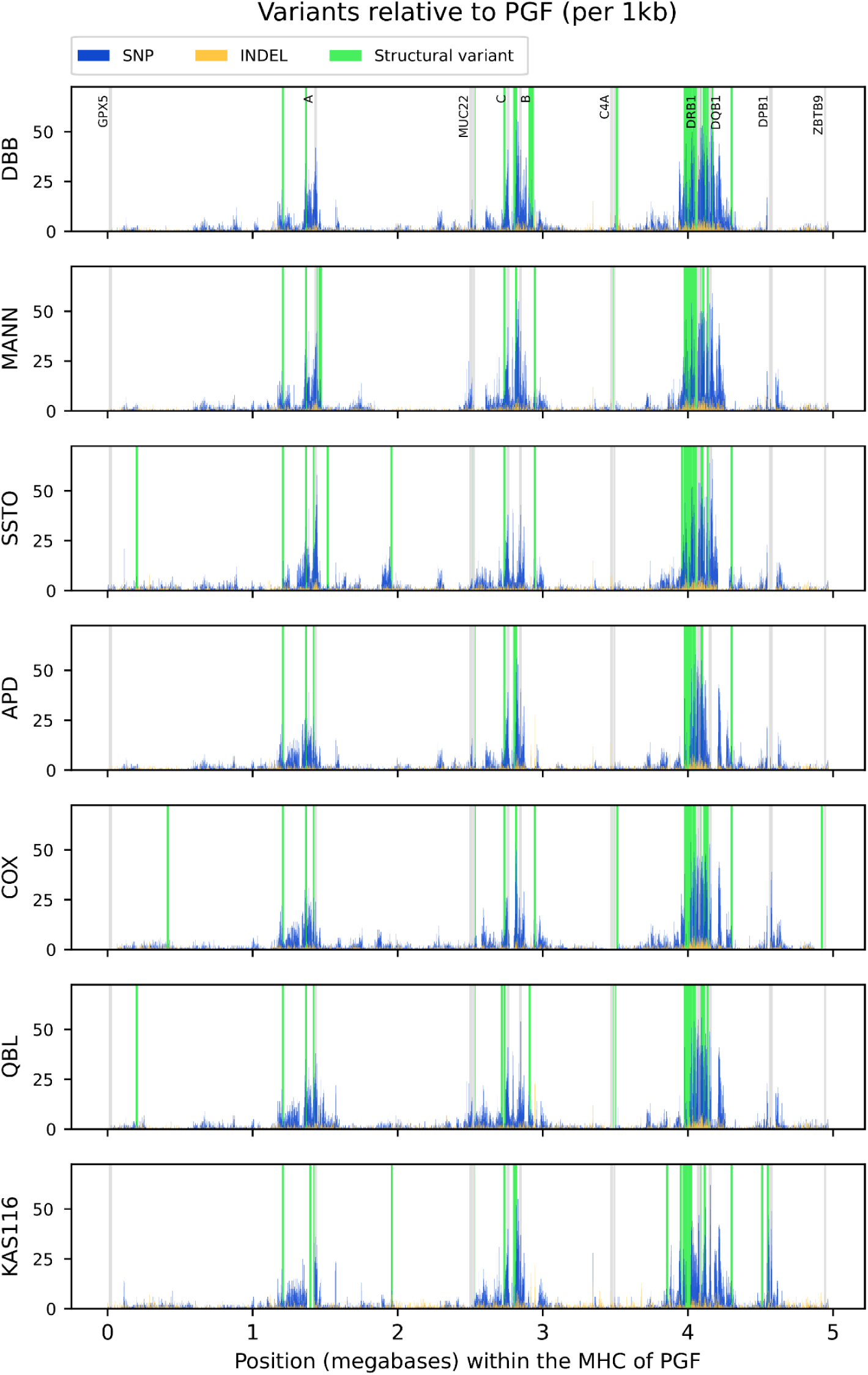
Sequence and structural variation of the completed *MHC* haplotypes. Shown are the genomic variations relative to the GRCh38 reference (PGF) within each completed *MHC* region haplotype. (blue) single-nucleotide polymorphism (SNP), (gold) insertion-deletion (INDEL), (green) structural variants (indels > 1000bp, or inversions). (grey) selected genes for orientation (names indicated at top). The variants were counted in 1kbp non-overlapping windows. See the Methods section for details.

### Analysis of *MHC class II* structures

The *class II* region is the most structurally variable region of the human *MHC*; it is characterized by four major haplotype structures that are defined by the genotype of the *HLA-DRB1* gene and *HLA-DRB3, -4* and *-5* carrier status. In accordance with Trowsdale *et al*. ^20^, we refer to the four major categories of *MHC class II* structure as DR1 (*DRB1*01* or *\*10*, with no *DRB3-5*), DR2 (*DRB1*15* or *\*16*, with *DRB5*), DR3 (*DRB1*03, *11, *12, *13* or *\*14* with *DRB3*), DR4 (*DRB1*04, *09* or *\*07* with *DRB4*). DR8 (*DRB1*08*, with no *DRB3-*5), which was not targeted here, is distinguished from DR1 by lacking the *DRB*6 pseudogene.

When compared to GRCh38, the haplotypes presented here represent the first complete assemblies of DR1 (*DRB1* only) and DR4 (*DRB1* + *DRB4*) *MHC class II* sequence structures. For the following analysis, the *MHC class II* region was defined as the region from *HLA-DRA* to 20 kb downstream of *HLA-DRB1*, and results are reported separately for the DRB3/4/5 carrier status-defined haplotype classes.

#### DR3 (*DRB1* + *DRB3*) haplotypes

The DR3 haplotype structure is carried by the APD, COX, and QBL cell lines. The assembled *MHC class II* haplotypes of APD and QBL are 149.5 and 149.4 kbp in length, respectively, compared to the previously characterized 153.4 kbp for the haplotype from COX. Both multiple sequence and pairwise alignments confirmed the structural homology of the assembled *DRB3* containing haplotypes (Figure 3, Supplementary Figure 3). Of note, the small difference in *MHC class II* haplotype length within the DR3 group is largely attributable to a ∼3.5 kbp sequence segment carried by COX (approximate position: 38 kbp downstream of *HLA-DRA*; Figure 4, blue arrow) that is shared with the DR4 group and KAS116 (DR1) haplotypes, but absent from the other DR3 haplotypes. We also identified a ∼4kb inversion (relative to the DR4 group and KAS116 haplotypes) carried by the DR3 group and PGF (DR2) haplotypes, located approximately 80 kbp and 60 kbp downstream of *HLA-DRA* in COX and (in its reverse-complemented form) in SSTO, respectively. The existence of this inversion was confirmed with minimap2^43^. Interspersed repeats accounted for 56 – 57% of sequence content (Supplementary Table 5) in the *MHC class II* region of the DR3 group; the positions of repeat elements are conserved across the three DR3 haplotypes (Figure 4).

**Figure 3:**
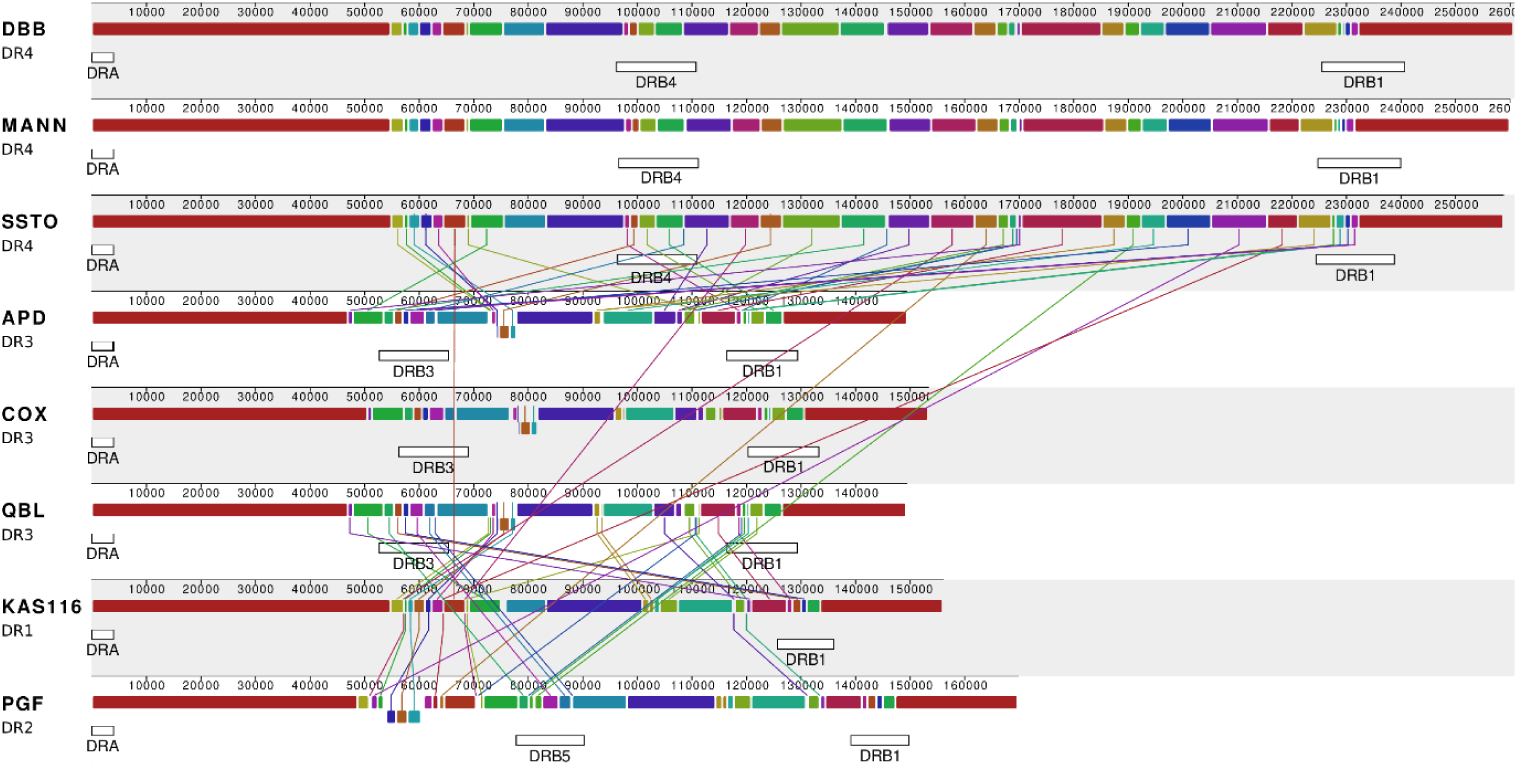
Multiple-sequence alignment visualization of *MHC class II* haplotype structures. Shown is a comparison of the eight completed *MHC class II* region sequences. Colours represent respective sequence similarity across haplotypes. Segments drawn underneath the respective plots represent inversions. The plot was created using Mauve ^51^ with parameter ‘seed weight’ set to 22. Vertical lines connecting horizontally aligned homologous regions were edited for clarity.

**Figure 4:**
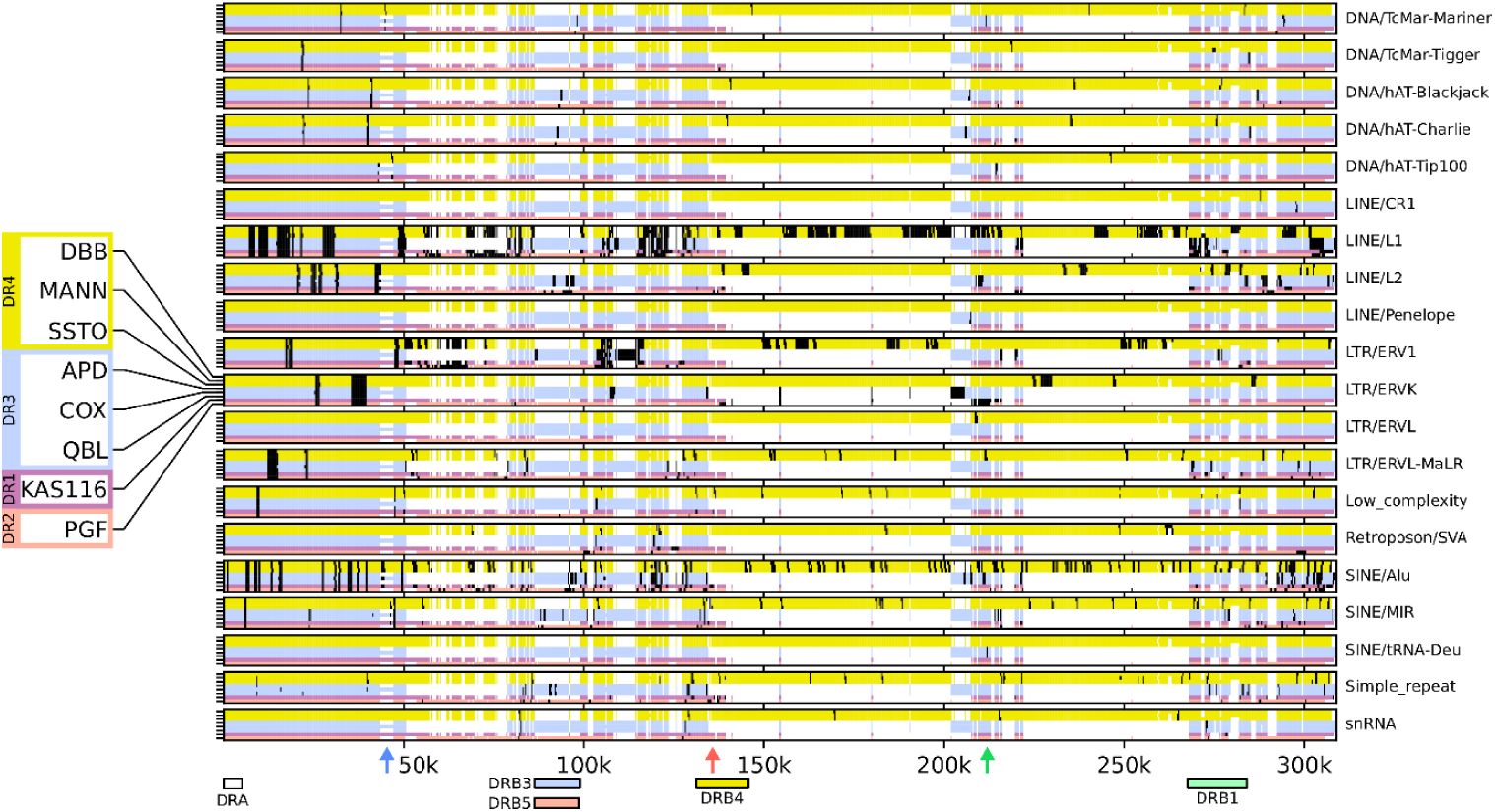
Repeat elements within the *MHC class II* region. Shown are positions of repeat elements identified in the *MHC class II* region of each of the eight completed haplotypes. The haplotypes are coloured by their broad grouping: (yellow) DR4, (blue) DR3, (purple) DR1, (red) DR2. White indicates gaps relative to the longest sequence alignment. Black shading indicates locations where the respective repeat element was identified. The multiple sequence alignment was generated using mafft ^56^. Repeat elements were identified using RepeatMasker ^55^, where the sequences were divided into 300bp non-overlapping windows and those windows having any overlap with a repeat element were counted as containing a repeat. Red and green arrows indicate the positions of *HLA-DRB6* and ERV that are components of the segment distinguishing DR1 from DR4; a blue arrow, the position of the identified small structural variant that is not in strict LD with any specific *MHC class II* haplotype.

#### DR4 (*DRB1* + *DRB4*) haplotypes

The DR4 haplotype structure is carried by the DBB, MANN and SSTO cell lines. The assembled *MHC class II* haplotypes of the DR4 group range in length from 258.8 - 260.8 kbp; of note, GRCh38 does not include a fully resolved *MHC class II* haplotype of the DR4 group. We used both multiple sequence and pairwise alignment to investigate the extent of structural homology within the DR4 group and found no large structural variants (Figure 3; Supplementary Figure 3). Repeat element positions are highly conserved across the DR4 haplotypes. The exceptions are retrotransposon/SVA and LINE1 insertions upstream of *HLA-DRB1* in DBB and SSTO, and LINE1 insertions in the *HLA-DRB1* of DBB and MANN (Figure 4), which characterize minor structural differences between the haplotypes. At ∼63%, the interspersed repeat content of the DR4 group is higher than that of any other group (Supplementary Table 5). Repeat elements also account for a considerable fraction of the sequence that distinguishes the DR4 group from other *MHC class II* sequences (Figure 4).

#### DR2 (*DRB1* + *DRB5*) haplotypes

The DR2 haplotype structure is carried by PGF, which was used for the canonical chromosome 6 reference *MHC* haplotype of GRCh38. The *MHC class II* region of PGF has a length of 169.9 kb; as PGF is the only representative of the DR2 group, no within-group structural variation analysis was carried out. Interspersed repeats account for 62.6% of the *MHC class II* sequence content of PGF (Supplementary Table 5). High density of repeat elements (in particular LINE/L1 and LTR/ERV1) is also found in the *class II* sequence stretches of PGF, and do not align to any of the other assembled haplotypes (Figure 4).

#### DR1 (*DRB1* only) haplotype

The DR1 haplotype structure is carried by KAS116, whereas GRCh38 contains no representative of this haplotype structure. The *MHC class II* haplotype of KAS116 has a length of 156.1 kbp. As KAS116 is the only representative of the DR1 group, no within-group structural variation analysis could be carried out. Multiple and pairwise sequence alignments were computed to compare the haplotype of KAS116 to the other *MHC class II* haplotype groups (Figure 3; Supplementary Figure 3). The DR1 is most similar to the DR4 haplotypes, where KAS116 differs by replacement of an approximately 120 kbp segment between *HLA-DRB4* an *HLA-DRB1* with an approximately 20 kbp segment (Supplementary Figure 3). The segment includes a copy of *HLA-DRB6* at its 3’ end, followed by a 7.5kb long ERV sequence (Figure 4, red and green arrows). Interspersed repeats account for 61.5% of the *MHC class II* sequence content of KAS116 (Supplementary Table 5).

### Improvement of short-read mapping experiment with newly assembled *MHC* haplotypes

To investigate the potential benefits for mapping whole-genome short-read sequencing data conferred by using the fully resolved *MHC* haplotypes presented here instead of the incomplete versions currently part of GRCh38, we performed a comparative mapping experiment. Whole-genome Illumina sequencing reads from seven samples (two samples homozygous for a PGF-like *MHC class II* structure; two samples homozygous for an APD-like *MHC class II* structure; and three samples homozygous for the DR1 *MHC class II* structure) were aligned to a standard version of the GRCh38 reference genome (containing incompletely resolved *MHC* ‘alt_ref’ contigs) and to an improved version of GRCh38 containing the complete assemblies presented here added (see Methods). We observed only a small increase in the total (whole-genome) number of mapped reads due to the inclusion of the improved *MHC* reference sequences (Table 2). However, the number of reads recruited to the full-length *MHC* contigs as part of ‘proper’ read pairs (that is, with correct read orientations and with an insert size deemed plausible by the short-read mapper BWA) increased by 0.32% – 0.69% (Table 2). The largest effects were observed for sample NA10847 (0.95% increase), which carries a DR1 *MHC class II* structure (not represented in GRCh38), and for sample NA20861 (0.51%), which carries an APD-like *MHC class II* structure (representing the least-complete GRCh38 *MHC* ‘alt_ref’ contig; 2.5 Mbp missing bases; Supplementary Table 4).

## Discussion

To improve the scope and utility of reference data for population-scale immunogenomics, we have assembled and resolved the structure and sequence of five *MHC* reference haplotypes. This work completed five targets of the ‘*MHC haplotype Project*’ ^37^ that currently form part of the human reference genome. In targeting the same cell lines present in the GRCh38 human reference genome build, we increased the number of fully resolved *MHC* haplotypes from two to seven. Importantly, we include the first fully resolved *MHC class II* sequences representing the DR4 *MHC* haplotype group, which is 55-76kbp longer than the PGF sequence that forms the baseline reference for GRCh38. In addition, our analysis of cell line KAS116 provides a structurally resolved reference haplotype for the DR1 *MHC class II* structure that is not yet represented in GRCh38; polishing this sequence with whole-genome Illumina sequencing data to remove residual INDEL errors remains as future work. When combined with the complete PGF (DR2) and COX (DR3) reference haplotypes from GRCh38, the sequences presented here provide a comprehensive reference panel covering the major *MHC class II* haplotype structures.

Despite progress in long-read sequencing technologies, and their combination with other technologies, fully resolving complex repeat structures of the *MHC*, in for example the *MHC class II* or *C4* regions, remains a challenge ^62^. Here we employed a hybrid strategy, integrating ultra-long Nanopore sequencing data, previously assembled, highly accurate short contigs from targeted sequencing, and Illumina whole-genome sequencing data. For assembly we devised custom bioinformatics pipelines, where the long-read data provided structural integrity, the contigs provided sequence accuracy, and the whole-genome reads were vital for haplotype polishing. The accuracy of the assembled sequences is supported by multiple lines of evidence, including (i) structural consistency between available GRCh38 assembly fragments and the assemblies we produced; (ii) internal consistency within the different *MHC class II* haplotype groups; and (iii) comparable rates of genetic variation relative to the canonical PGF haplotype between the haplotypes assembled here and the other complete *MHC* reference haplotype part of GRCh38, COX (Table 1b). Of note, we observed higher INDEL rate for the *MHC* assembly of cell line KAS116; this may reflect the increased INDEL error rate of Nanopore-based assembly consensus sequences in combination with the fact that polishing based on whole-genome Illumina sequencing data was not performed for KAS116. Future goals include complete polishing of this DR1 reference, as well as targeting representative DR8 haplotypes, which also lack the *DRB3, 4* or *5* genes, to determine the ancestral relationship with DR1.

The availability of a comprehensive set of haplotype sequences enabled us to assess variation across the major *MHC class II* structures. Specifically, we ruled out the presence of any previously uncharacterized large-scale structural variants, and showed that the DR1 *class II* haplotype structure is most closely related to that of the DR4 haplotype. Interestingly, we also identified a small structural variant in the *class II* region that is not in strict LD with any specific *MHC class II* haplotype class (Figure 4, blue arrow). Although we did not carry out a full analysis of the assembled *MHC class II* sequences, we have demonstrated that the density of repeat elements increases in the *MHC class II* region compared to the rest of the *MHC* (Supplementary Table 5), that it varies between *MHC class II* haplotype groups, and that repeat elements are often found in the group-exclusive sequence regions that differentiate between the *class II* haplotype groups. The latter observation suggests a role for the repeat elements in the divergence and / or maintenance of separation between the different haplotypes^21^. Our findings confirm and expand recent detailed analyses of the *MHC* haplotypes ^21,63^.

As a first step towards measuring how improved *MHC* reference assemblies can contribute to improved read mapping, and sequence and structure variant determination throughout the *MHC* genomic region, we performed a comparative short-read mapping experiment. This experiment showed that our improved assemblies enable improvements in the recruitment of ‘proper pair’ read alignments to full-length *MHC* reference contigs. As expected, the largest effects were observed for the samples with *MHC class II* structures that were unfinished in GRCh38. Further improvements are expected by increasing the representation of *MHC* haplotypes of non-European ancestry, which could complement ongoing pan-genomic assembly efforts ^64,65^, and by the further development of graph-based approaches integrating information across reference sequences and haplotypes ^27,33,34,66-69^. Efforts along both directions are currently under way and we predict significant improvements to accessibility of immunogenetic variation and its phenotypic impacts in the near future.

**Supplementary Table 1: Studied cell lines and their HLA genotypes**. Shows the *HLA class I* and *II* genotypes of the six MHC homozygous cell lines analysed here, and two that were completed previously (PGF and COX). (NB: MANN is also known as MOU).

**Supplementary Table 2: Data and assembly generation summary**. Illumina and Nanopore sequencing data summary for the sequenced cell lines, number of polishing rounds performed during the assembly process (see Methods), and BioSample and GenBank IDs.

**Supplementary Table 3: Data generation details**. Illumina and Nanopore sequencing runs performed on the different cell lines, including details on the utilized DNA extraction, library preparation and sequencing protocols.

**Supplementary Table 4: Assembly comparison to GRCh38 and Norman et al. (2017)**. Comparison between the haplotype assemblies generated here and earlier assemblies of the same haplotypes from GRCh38 and the scaffold dataset released by ^9^. “Undetermined nucleotides” were quantified by counting the number of ‘N’ characters in the corresponding assembly version; “missing bases” were quantified by mapping the earlier assemblies of the considered haplotypes against the versions presented here, retaining only unique alignments, and counting the number of bases with 0 coverage from the earlier-assembly alignments.

**Supplementary Table 5: Interspersed repeats content**. Proportion of assembled *MHC* sequences marked as interspersed repeats by the RepeatMasker ^55^ algorithm, reported separately for the complete *MHC* and the *MHC class II* subregion.

TODO delete ! ^9^ (S^51^

**Supplementary Figure 1: C4 genotypes of assembled *MHC* sequences and the complete GF and COX *MHC* reference haplotypes from GRCh38**. HERV = human endogenous retrovirus. Visual display adapted from Sekar *et al*. ^26^.

**Supplementary Figure 2: Mauve multiple-sequence alignment of the assembled *MHC* sequences and the complete PGF and COX *MHC* reference haplotypes from GRCh38**. The plot is based on an alignment generated with the ‘seed weight’ parameter set to 22.

**Supplementary Figure 3: Dot plots visualizing the haplotype structures of the assembled *MHC* sequences and the complete PGF and COX *MHC* reference haplotypes from GRCh38**. Each plot consists of two triangular panels, showing visualizations of whole-*MHC* (upper-left half) and *MHC class II* (lower-right half) structures of the corresponding haplotypes (see labels on the X and Y axes, also showing the lengths of the visualized sequences. Panel (a) shows a comparison between one representative of each of the four major *MHC class II* haplotype structures; panel (b) shows a comparison between the sequence structures of the DR3 group of *MHC* haplotypes; panel (c), a comparison between the sequence structures of the DR4 *MHC* haplotype group. Shaded bars in the *MHC class II*-specific panels indicate the positions of the *HLA-DRB3, -DRB4*, and *-DRB5* genes. The plots were generated with the nucmer (parameters --maxmatch --nosimplify --mincluster 300), mummerplot ^46^ and gnuplot programs.

## Acknowledgements

This work was supported by the National Institutes of Health of the USA (NIAID U01 AI090905); the Forschungskommission of the Medical Faculty of Heinrich Heine University Düsseldorf; the Jürgen Manchot Foundation; the German Federal Ministry of Education and Research (Bundesministerium für Bildung und Forschung; award numbers 031L0184B); the German Research Foundation (award 428994620). We would like to thank Tamim Shaikh and Elizabeth Geiger for help with the PromethIon machine.

Computational support and infrastructure were provided by the “Centre for Information and Media Technology” (ZIM) at the University of Düsseldorf (Germany).

## Data and Code Availability

Whole-genome Nanopore and Illumina sequencing data generated as part of this study and the generated *MHC* assemblies were submitted to NCBI BioProject PRJNA764575. The whole-genome sequencing data were filtered to contain only reads mapping to the assembled *MHC* sequences prior to submission. The generated assemblies were also submitted to GenBank; GenBank accessions are listed in Table 1a. Data generated by ^9^ that were used as part of the assembly process described here are also publicly available; the corresponding BioSample IDs are listed in Supplementary Table 2.

The Python source code used for annotation of the assembled *MHC* sequences is available via PyPI (https://pypi.org/project/MHC-Annotation/). A separate publication describing the annotation method is currently under preparation.

## References

1. Trowsdale J, Knight JC. Major histocompatibility complex genomics and human disease. Annu Rev Genomics Hum Genet. 2013;14:301–323.

2. Matzaraki V, Kumar V, Wijmenga C, Zhernakova A. The MHC locus and genetic susceptibility to autoimmune and infectious diseases. Genome Biol. 2017;18(1):76.

3. Dendrou CA, Petersen J, Rossjohn J, Fugger L. HLA variation and disease. Nature reviews Immunology. 2018;18(5):325–339.

4. Petersdorf EW. Role of major histocompatibility complex variation in graft-versus-host disease after hematopoietic cell transplantation. F1000Res. 2017;6:617.

5. Horton R, Wilming L, Rand V, et al. Gene map of the extended human MHC. Nat Rev Genet. 2004;5(12):889–899.

6. Kellis M, Wold B, Snyder MP, et al. Defining functional DNA elements in the human genome. Proc Natl Acad Sci U S A. 2014;111(17):6131–6138.

7. Apps R, Qi Y, Carlson JM, et al. Influence of HLA-C expression level on HIV control. Science. 2013;340(6128):87–91.

8. Jin Y, Gittelman RM, Lu Y, et al. Evolution of DNAase I Hypersensitive Sites in MHC Regulatory Regions of Primates. Genetics. 2018;209(2):579–589.

9. Norman PJ, Norberg SJ, Guethlein LA, et al. Sequences of 95 human MHC haplotypes reveal extreme coding variation in genes other than highly polymorphic HLA class I and II. Genome Res. 2017;27(5):813–823.

10. Degli-Esposti MA, Leaver AL, Christiansen FT, Witt CS, Abraham LJ, Dawkins RL. Ancestral haplotypes: conserved population MHC haplotypes. Hum Immunol. 1992;34(4):242–252.

11. Ahmad T, Neville M, Marshall SE, et al. Haplotype-specific linkage disequilibrium patterns define the genetic topography of the human MHC. Hum Mol Genet. 2003;12(6):647–656.

12. Cullen M, Perfetto SP, Klitz W, Nelson G, Carrington M. High-resolution patterns of meiotic recombination across the human major histocompatibility complex. Am J Hum Genet. 2002;71(4):759–776.

13. Parham P, Ohta T. Population biology of antigen presentation by MHC class I molecules. Science. 1996;272(5258):67–74.

14. Deng Z, Zhen J, Harrison GF, et al. Adaptive Admixture of HLA Class I Allotypes Enhanced Genetically Determined Strength of Natural Killer Cells in East Asians. Mol Biol Evol. 2021;38(6):2582–2596.

15. Dilthey AT. State-of-the-art genome inference in the human MHC. Int J Biochem Cell Biol. 2021;131:105882.

16. de Bakker PI, McVean G, Sabeti PC, et al. A high-resolution HLA and SNP haplotype map for disease association studies in the extended human MHC. Nat Genet. 2006;38(10):1166–1172.

17. Yunis EJ, Larsen CE, Fernandez-Viña M, et al. Inheritable variable sizes of DNA stretches in the human MHC: conserved extended haplotypes and their fragments or blocks. Tissue Antigens. 2003;62(1):1–20.

18. Church DM. Genomes for all. Nat Biotechnol. 2018;36(9):815–816.

19. Kennedy AE, Ozbek U, Dorak MT. What has GWAS done for HLA and disease associations? Int J Immunogenet. 2017;44(5):195–211.

20. Trowsdale J, Young JA, Kelly AP, et al. Structure, sequence and polymorphism in the HLA-D region. Immunol Rev. 1985;85:5–43.

21. Kulski JK, Suzuki S, Shiina T. Haplotype Shuffling and Dimorphic Transposable Elements in the Human Extended Major Histocompatibility Complex Class II Region. Front Genet. 2021;12:665899.

22. Jeffreys AJ, Kauppi L, Neumann R. Intensely punctate meiotic recombination in the class II region of the major histocompatibility complex. Nat Genet. 2001;29(2):217–222.

23. Wu YL, Savelli SL, Yang Y, et al. Sensitive and specific real-time polymerase chain reaction assays to accurately determine copy number variations (CNVs) of human complement C4A, C4B, C4-long, C4-short, and RCCX modules: elucidation of C4 CNVs in 50 consanguineous subjects with defined HLA genotypes. J Immunol. 2007;179(5):3012–3025.

24. Moutsianas L, Jostins L, Beecham AH, et al. Class II HLA interactions modulate genetic risk for multiple sclerosis. Nat Genet. 2015;47(10):1107–1113.

25. Pappas DJ, Lizee A, Paunic V, et al. Significant variation between SNP-based HLA imputations in diverse populations: the last mile is the hardest. Pharmacogenomics J. 2018;18(3):367–376.

26. Sekar A, Bialas AR, de Rivera H, et al. Schizophrenia risk from complex variation of complement component 4. Nature. 2016;530(7589):177–183.

27. Dilthey A, Cox C, Iqbal Z, Nelson MR, McVean G. Improved genome inference in the MHC using a population reference graph. Nat Genet. 2015;47(6):682–688.

28. Dilthey AT, Mentzer AJ, Carapito R, et al. HLA*LA-HLA typing from linearly projected graph alignments. Bioinformatics. 2019;35(21):4394–4396.

29. Dilthey AT, Gourraud PA, Mentzer AJ, Cereb N, Iqbal Z, McVean G. High-Accuracy HLA Type Inference from Whole-Genome Sequencing Data Using Population Reference Graphs. PLoS Comput Biol. 2016;12(10):e1005151.

30. Xie C, Yeo ZX, Wong M, et al. Fast and accurate HLA typing from short-read next-generation sequence data with xHLA. Proc Natl Acad Sci U S A. 2017;114(30):8059–8064.

31. Lee H, Kingsford C. Kourami: graph-guided assembly for novel human leukocyte antigen allele discovery. Genome Biol. 2018;19(1):16.

32. Kim D, Paggi JM, Park C, Bennett C, Salzberg SL. Graph-based genome alignment and genotyping with HISAT2 and HISAT-genotype. Nat Biotechnol. 2019;37(8):907–915.

33. Eggertsson HP, Kristmundsdottir S, Beyter D, et al. GraphTyper2 enables population-scale genotyping of structural variation using pangenome graphs. Nat Commun. 2019;10(1):5402.

34. Eggertsson HP, Jonsson H, Kristmundsdottir S, et al. Graphtyper enables population-scale genotyping using pangenome graphs. Nat Genet. 2017;49(11):1654–1660.

35. Horton R, Gibson R, Coggill P, et al. Variation analysis and gene annotation of eight MHC haplotypes: the MHC Haplotype Project. Immunogenetics. 2008;60(1):1–18.

36. Traherne JA, Horton R, Roberts AN, et al. Genetic analysis of completely sequenced disease-associated MHC haplotypes identifies shuffling of segments in recent human history. PLoS Genet. 2006;2(1):e9.

37. Stewart CA, Horton R, Allcock RJ, et al. Complete MHC haplotype sequencing for common disease gene mapping. Genome Res. 2004;14(6):1176–1187.

38. Allcock RJ, Atrazhev AM, Beck S, et al. The MHC haplotype project: a resource for HLA-linked association studies. Tissue Antigens. 2002;59(6):520–521.

39. Schneider VA, Graves-Lindsay T, Howe K, et al. Evaluation of GRCh38 and de novo haploid genome assemblies demonstrates the enduring quality of the reference assembly. Genome Res. 2017;27(5):849–864.

40. Norman PJ, Norberg SJ, Nemat-Gorgani N, et al. Very long haplotype tracts characterized at high resolution from HLA homozygous cell lines. Immunogenetics. 2015;67(9):479–485.

41. Mickelson E, Hurley C, Ng J, et al. 13th IHWS Shared Resources Joint Report. IHWG Cell and Gene Bank and reference cell panels. In: Ja H, ed. Immunobiology of the Human MHC: Proceedings of the 13th International Histocompatibilty Workshop and Conference. Vol 1. Seattle: IHWG Press; 2006:523–553.

42. Jain M, Koren S, Miga KH, et al. Nanopore sequencing and assembly of a human genome with ultra-long reads. Nat Biotechnol. 2018;36(4):338–345.

43. Li H. Minimap2: pairwise alignment for nucleotide sequences. Bioinformatics. 2018;34(18):3094–3100.

44. Koren S, Walenz BP, Berlin K, Miller JR, Bergman NH, Phillippy AM. Canu: scalable and accurate long-read assembly via adaptive k-mer weighting and repeat separation. Genome Res. 2017;27(5):722–736.

45. Kolmogorov M, Yuan J, Lin Y, Pevzner PA. Assembly of long, error-prone reads using repeat graphs. Nat Biotechnol. 2019;37(5):540–546.

46. Kurtz S, Phillippy A, Delcher AL, et al. Versatile and open software for comparing large genomes. Genome Biol. 2004;5(2):R12.

47. Li H. Aligning sequence reads, clone sequences and assembly contigs with BWA-MEM. 2013: 1303.3997. https://ui.adsabs.harvard.edu/abs/2013arXiv1303.3997L. Accessed March 01, 2013.

48. DePristo MA, Banks E, Poplin R, et al. A framework for variation discovery and genotyping using next-generation DNA sequencing data. Nat Genet. 2011;43(5):491–498.

49. Robinson JT, Thorvaldsdottir H, Winckler W, et al. Integrative genomics viewer. Nat Biotechnol. 2011;29(1):24–26.

50. Li H, Handsaker B, Wysoker A, et al. The Sequence Alignment/Map format and SAMtools. Bioinformatics. 2009;25(16):2078–2079.

51. Darling AC, Mau B, Blattner FR, Perna NT. Mauve: multiple alignment of conserved genomic sequence with rearrangements. Genome Res. 2004;14(7):1394–1403.

52. Robinson J, Barker DJ, Georgiou X, Cooper MA, Flicek P, Marsh SGE. IPD-IMGT/HLA Database. Nucleic Acids Res. 2020;48(D1):D948–D955.

53. O’Leary NA, Wright MW, Brister JR, et al. Reference sequence (RefSeq) database at NCBI: current status, taxonomic expansion, and functional annotation. Nucleic Acids Res. 2016;44(D1):D733–745.

54. Yang Y, Chung EK, Wu YL, et al. Gene copy-number variation and associated polymorphisms of complement component C4 in human systemic lupus erythematosus (SLE): low copy number is a risk factor for and high copy number is a protective factor against SLE susceptibility in European Americans. Am J Hum Genet. 2007;80(6):1037–1054.

55. Smit AFA, Hubley R, Green P. RepeatMasker Open-4.0. http://www.repeatmasker.org. Published 2013-2015. Accessed.

56. Katoh K, Standley DM. MAFFT multiple sequence alignment software version 7: improvements in performance and usability. Mol Biol Evol. 2013;30(4):772–780.

57. Byrska-Bishop M, Evani US, Zhao X, et al. High coverage whole genome sequencing of the expanded 1000 Genomes Project cohort including 602 trios. bioRxiv. 2021.

58. 1000 Genomes Project Consortium, Auton A, Brooks LD, et al. A global reference for human genetic variation. Nature. 2015;526(7571):68–74.

59. Abi-Rached L, Gouret P, Yeh JH, et al. Immune diversity sheds light on missing variation in worldwide genetic diversity panels. PLoS One. 2018;13(10):e0206512.

60. Gourraud PA, Khankhanian P, Cereb N, et al. HLA diversity in the 1000 genomes dataset. PLoS One. 2014;9(7):e97282.

61. Smith WP, Vu Q, Li SS, Hansen JA, Zhao LP, Geraghty DE. Toward understanding MHC disease associations: partial resequencing of 46 distinct HLA haplotypes. Genomics. 2006;87(5):561–571.

62. Chin CS, Wagner J, Zeng Q, et al. A diploid assembly-based benchmark for variants in the major histocompatibility complex. Nat Commun. 2020;11(1):4794.

63. Kulski JK, Suzuki S, Shiina T. SNP-Density Crossover Maps of Polymorphic Transposable Elements and HLA Genes Within MHC Class I Haplotype Blocks and Junction. Front Genet. 2020;11:594318.

64. Ebert P, Audano PA, Zhu Q, et al. Haplotype-resolved diverse human genomes and integrated analysis of structural variation. Science. 2021;372(6537).

65. Miga KH, Wang T. The Need for a Human Pangenome Reference Sequence. Annu Rev Genomics Hum Genet. 2021;22:81–102.

66. Rakocevic G, Semenyuk V, Lee WP, et al. Fast and accurate genomic analyses using genome graphs. Nat Genet. 2019;51(2):354–362.

67. Biederstedt E, Oliver JC, Hansen NF, et al. NovoGraph: Human genome graph construction from multiple long-read de novo assemblies. F1000Res. 2018;7:1391.

68. Garrison E, Siren J, Novak AM, et al. Variation graph toolkit improves read mapping by representing genetic variation in the reference. Nat Biotechnol. 2018;36(9):875–879.

69. Ebler J, Clarke WE, Rausch T, et al. Pangenome-based genome inference. bioRxiv. 2020.

