## Supplementary figures and images for "Complete sequences of six Major Histocompatibility Complex haplotypes, including all the major *MHC class II* structures"

### Supplementary Figure 1

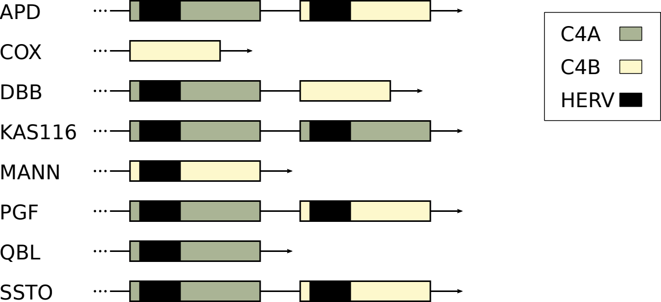

### Supplementary Figure 2

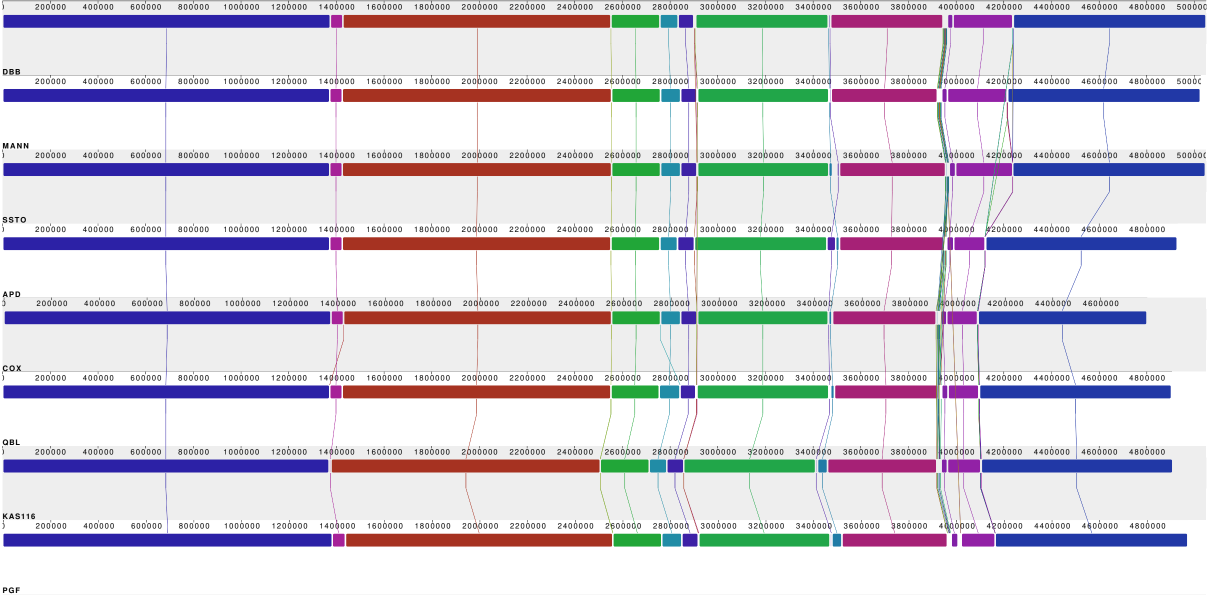

### Supplementary Figure 3

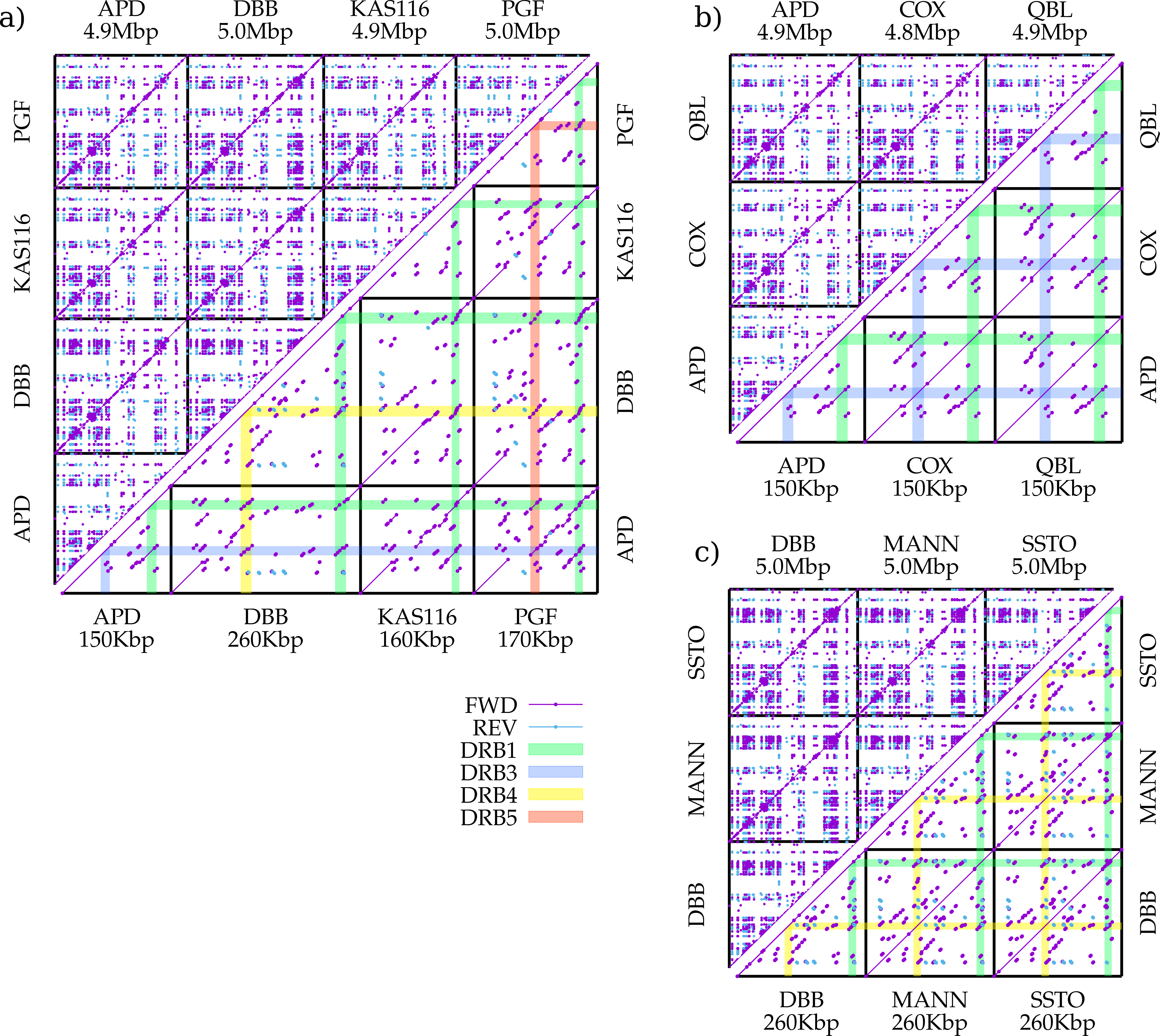
